# Hotwiring integrin endocytosis acutely modulates cell interactions

**DOI:** 10.1101/2024.06.24.600360

**Authors:** Sahil Kamboj, Alphonse Boché, Anneline Moret, Zixu Wang, Carole Aimé, Rémy Agniel, Johanne Leroy-Dudal, Franck Carreiras, Olivier Gallet, Stephen J Royle, Ambroise Lambert

## Abstract

Integrins are heterodimeric cell surface receptors that govern cell-cell interactions, which in turn can influence multiscale processes: cell migration, extracellular matrix remodeling and tissue formation. These processes occur over timescales which range from milliseconds to days. While various strategies exist to study integrin function across biological scales from cell to tissue, they are often chronic and fail to target specific cell-cell interactions acutely. We engineered cells to rapidly alter cell behavior by downregulating the surface population of α5β1 integrins through hot-wired clathrin-mediated endocytosis. This method allows for inducible, specific internalization of α5β1 integrins, achieving acute downregulation across various cell lines in 5-30 minutes. We show that induced internalization of α5β1 decreases the cell area, causes uptake of extracellular fibronectin, and decreases the rate of tumor spheroid compaction. This targeted control of multiscale processes by rapid downregulation of this important class of cell surface receptors demonstrates that hot-wired endocytosis is a useful tool to acutely modulate cell biology.

## Introduction

Integrins are cell surface receptors that mediate a wide array of cellular functions, including adhesion, migration, and signal transduction^1–4^. Despite the recognized importance of integrins in cancer and other biological contexts, studying their interactions and functional implications across different biological scales remains challenging. Integrins are heterodimeric proteins composed of α and β subunits^2,4^, which can bind extracellular matrix (ECM) proteins. This intricate relationship between integrins and the ECM is fundamental not only to physiological cellular and developmental processes but also in pathologic developments of diseases, including cancer, where integrin signaling is often dysregulated^5,6^.

Ovarian cancer displays a specific complex microenvironment, including ascites: a chronical fluid build-up in the peritoneal cavity that is rich in diverse proteins including soluble ECM proteins. The interaction of integrins with these proteins influences tumor growth, metastasis, angiogenesis, chemoresistance, and epithelial-mesenchymal transition (EMT). In particular, α5β1 integrin^7–9^ specifically recognizes the RGD (arginine-glycine-aspartate) motif present in fibronectin, facilitating cell adhesion and spreading on fibronectin-rich substrates^10^. α5β1 on the cell surface binds to fibronectin in the ECM, leading to the formation of focal adhesion complexes through the clustering of integrins. This binding initiates conformational changes leading to integrin activation, and further internalization of bound fibronectin through clathrin-mediated endocytosis. The α5β1-Fibronectin interaction not only results in the clustering of integrin receptors, facilitating the formation of a fibrillar matrix and ECM remodeling. It also plays a crucial role in determining the cellular response to the microenvironment^11–13^, making the number of its units decisive. In addition, the interaction between α5β1 and fibronectin regulates spheroid formation and adhesion. Cell-cell interactions during the assembly of multicellular aggregates, or spheroids, involve three critical stages: cell aggregation, compaction, and maturation. Initially, cell aggregation significantly relies on the interaction between cell surface integrins, notably α5β1 in fibronectin-poor microenvironment. This interaction ensures cell-cell cohesion, and blockade of integrin β1 disrupts spheroid assembly^14–16^. Integrins generate mechanical forces essential for packing multicellular aggregates in tight spheroids^17–19^.

Recently, an inducible endocytosis system was developed allowing for the rapid removal of a cell surface protein. Briefly, the FKBP-rapamycin-FRB inducible heterodimerization system was shown to induce the formation of clathrin-coated vesicles (CCVs), effectively bypassing the conventional regulatory steps involved in clathrin-mediated endocytosis (CME) and “hot-wiring” it^20^. It works by attaching a clathrin-binding protein (AP2B1) to a cell surface protein using rapamycin-induced heterodimerization of FKBP and FRB, which is sufficient to induce coated vesicle formation. This system showed that proteins can be rapidly downregulated (on a timescale of minutes) and that ‘hot-wired’ internalized vesicles perform similarly to their natural counterparts. This technology raises the possibility of rapidly modulating integrin surface receptor number if the system can be adapted for this purpose. Such an approach would have clear advantages over chronic methods to downregulate integrins such as knockdown or knockout (24h to 48h observations), giving access to phenomena with short timescales and fast kinetics.

We set out to investigate the modulation of cell surface integrins across different biological scales using a hot-wired endocytosis approach. We show that α5β1 integrins can be specifically targeted for endocytosis and that it influences crucial α5β1-fibronectin interactions. Thus, we demonstrate acute tuning of multiscale processes within minutes, ranging from cell adhesion and matrix remodeling to 3D spheroid formation, using a single molecular tool.

## Results

### Hot-wired endocytosis of integrin α5β1

Previously, it was demonstrated that induced heterodimerization can be used to trigger the endocytosis of proteins at the plasma membrane, so called hotwired endocytosis. To adapt this system to the triggered endocytosis of integrin α5β1, we engineered an α5-mCherry-FRB construct to heterodimerize with integrin β1 and, together with FKBP-AP2B1-GFP, trigger controllable internalization of the integrin α5β1 heterodimer (Figure 1A). We chose three different cancer cell lines for diversity in phenotypes (OVCAR, SKOV3 and HeLa). Independently of cell line type, we observed the formation of punctate structures resembling clathrin-coated vesicles after rapamycin treatment, for cells co-expressing α5-mCherry-FRB with FKBP-AP2B1-GFP (200 nM, 30 min; Figure 1B). Whereas in cells co-expressing α5-mCherry-FRB with GFP-FKBP, no puncta were formed. This indicated that we had successfully hot-wired endocytosis of integrin α5 in three different cell lines. We next used an image analysis procedure to document the effective trafficking of surface integrins to intracellular compartments. This analysis showed that relocalization after hotwiring in SKOV3 cells was more towards the nucleus upon induction of integrin internalization than in OVCAR3 cells (Figure 1C).

**Figure 1.**
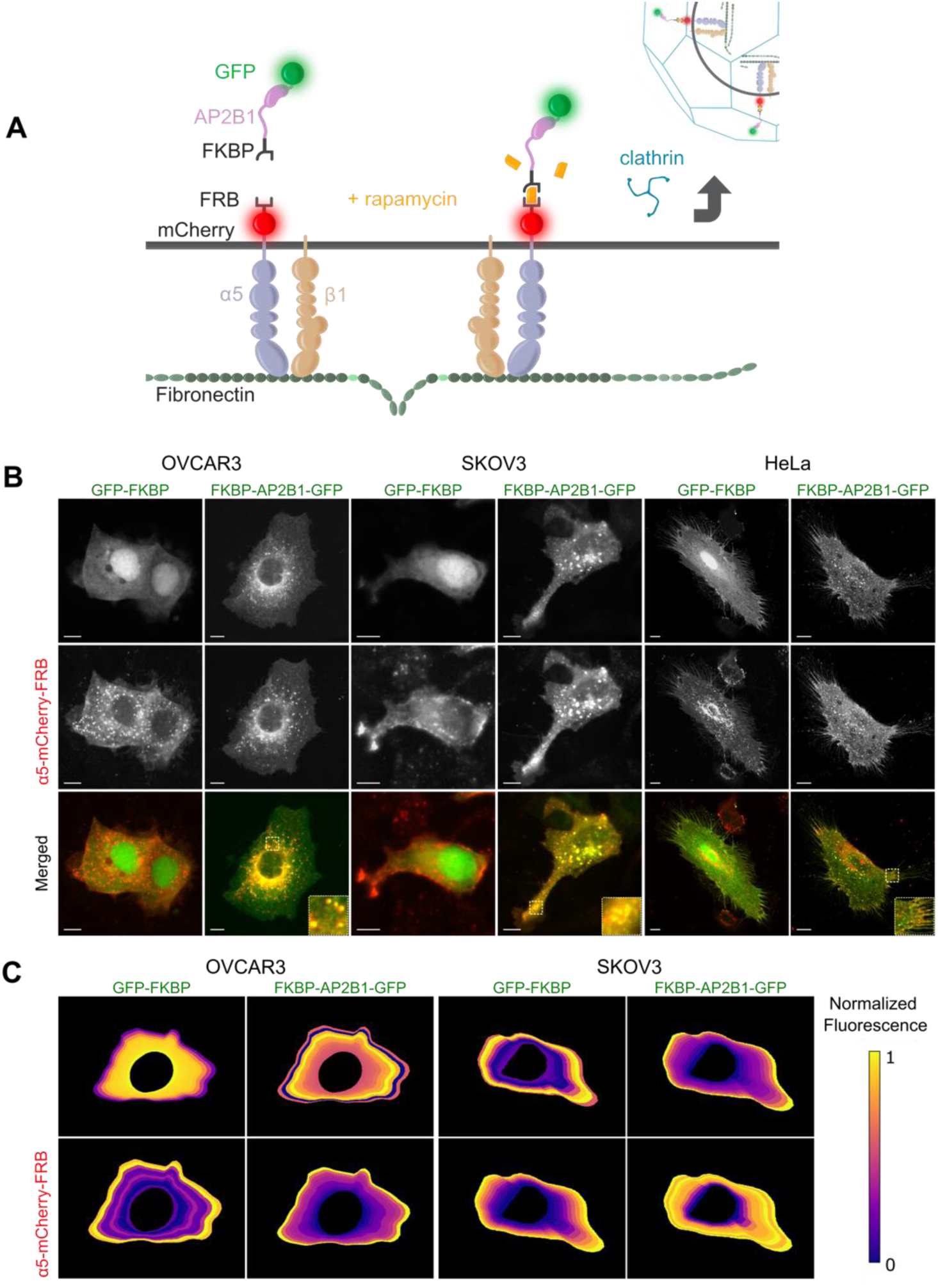
Hot-wired endocytosis of integrin α5. (A) Schematic diagram of hot-wired endocytosis of surface α5β1 integrin heterodimers. Cells co-express α5-mCherry-FRB and a clathrin hook (FKBP-AP2B1-GFP). Rapamycin induces the heterodimerization of FKBP and FRB domains meaning that the clathrin hook is recruited to α5 at the plasma membrane. Clathrin binds the clathrin hook and a clathrin-coated vesicle is formed containing α5β1 integrin heterodimers. (B) Representative micrographs of OVCAR3, SKOV3 or HeLa cells co-expressing α5-mCherry-FRB and either GFP-FKBP (control) or FKBP-AP2B1-GFP, and treated with rapamycin (200 nM, 30 min). Note the formation of colocalizing puncta in the FKBP-AP2B1-GFP but not GFP-FKBP conditions. Scale bar, 10 µm. (C) Distribution of integrins as a function of distance from nucleus to the membrane, mapped onto a typical cell profile. Normalized intensity per segment ((MeanValue-Minimum)/(Maximum-Minimum)) is shown for SKOV3 and OVCAR3 cells. N, >150 cells per condition in 3 independent experiments.

In order to confirm that our approach caused the internalization of expressed α5-mCherry-FRB and that this internalization represented α5β1 integrin heterodimers, we used an α5β1 antibody to label the surface population and understand if this signal was internalized. We could see robust uptake of surface labeled α5β1 integrin heterodimers by confocal microscopy (Supplementary Fig S2A). We also tested whether α5 that was internalized following induction was co-localized with early endosome antigen 1 (EEA-1, Supplementary Fig S2B). Indeed, we found colocalization of α5 puncta with EEA-1, 30 min post-induction but not in the GFP-FKBP control condition (Supplementary Fig S2).

Since integrin α5 and integrin αV both bind to fibronectin^20–23^, we investigated whether the hotwired endocytosis of α5 could inadvertently affect αV localization. Using immunofluorescence, we found no significant alteration in the distribution of αV following α5 internalization, which confirms the selectivity of our approach (Supplementary Fig S3).

Together these data indicate that we can hot-wire the endocytosis of integrin α5 in inducing the internalization of α5β1 integrin heterodimers specifically and target them to the endocytic pathway. This suggests that we can use this method as a tool to downregulate the surface population of α5β1 integrin heterodimers.

### Effect of induced internalization of α5 integrin on cell behavior

To investigate the effect and speed of hot-wired endocytosis of integrin α5, we performed live cell imaging in HeLa and SKOV3 cells co-expressing α5-mCherry-FRB and either FKBP-AP2B1-GFP or GFP-FKBP (control). Despite their similar morphologies, HeLa and SKOV3 cells exhibited significant differences in their response following induced internalization of integrin α5 upon rapamycin application (Figure 2). Colocalization of α5-mCherry-FRB and FKBP-AP2B1-GFP peaked at 50% in SKOV3 cells after 5 min of induction, while in HeLa cells, the peak was 40%, reached after 10 min of induction (Figure 2C,D).

**Figure 2.**
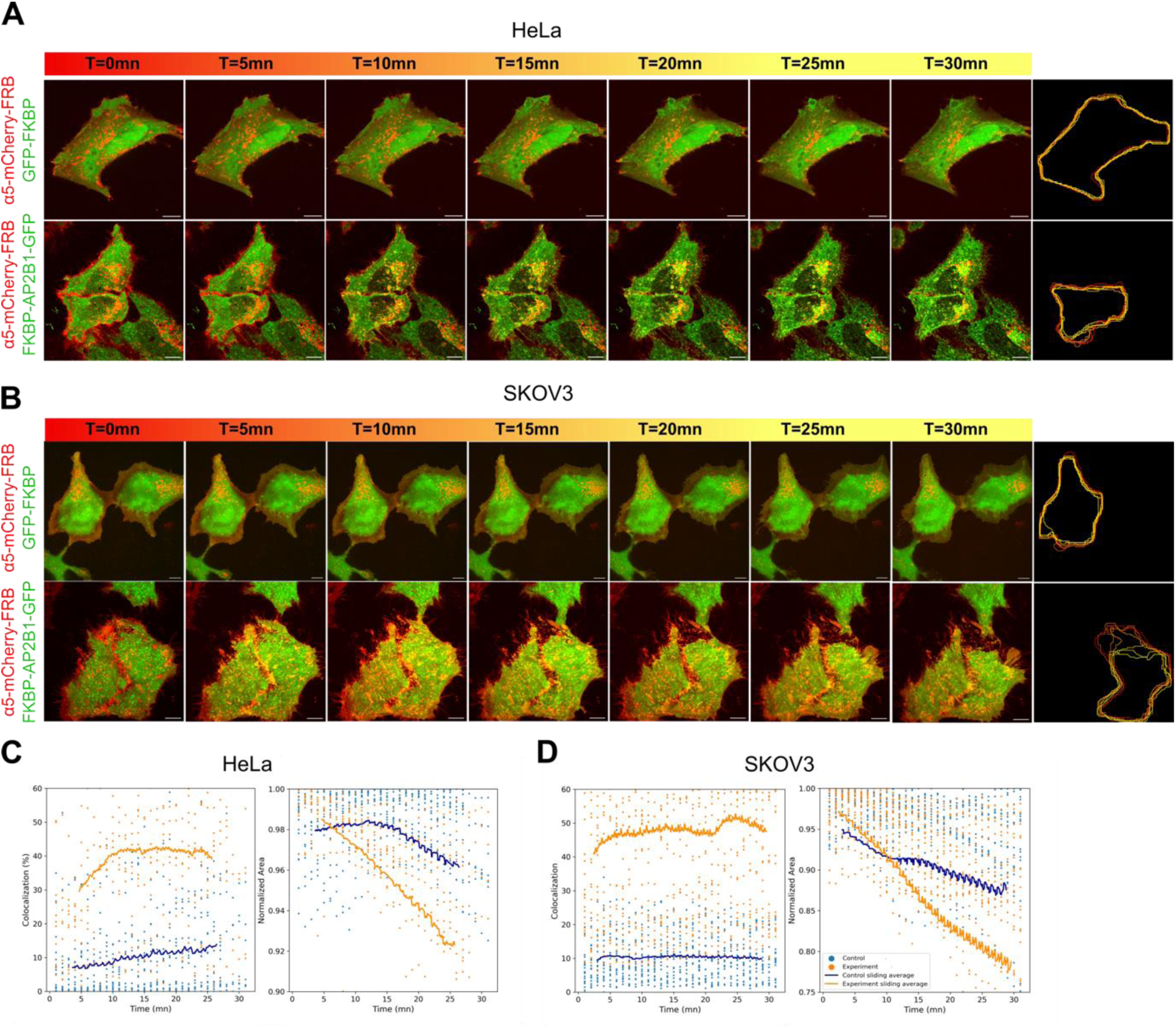
Induced internalization of integrin α5 causes a decrease in cell area. (A,B) Live cell imaging of HeLa (A) or SKOV3 (B) cells co-expressing α5-mCherry-FRB and either GFP-FKBP (control) or FKBP-AP2B1-GFP, treated with rapamycin (200 nM) at time 0. Right, overlaid cell profiles over time moving from red to yellow corresponding to cell profile over time. (C,D) Plots of colocalization of α5-mCherry-FRB with FKBP-AP2B1-GFP (orange) or GFP-FKBP (blue) (left) and normalized cell area (right) are shown for HeLa (C) and SKOV3 (D) cells. Dots indicate individual cell measurements; line is the average of all data. 27-30 cells were analyzed per condition per cell line over 30 mins in > 3 independent experiments. Scale bar, 10 µm

During these live cell imaging experiments, we noticed that the cell area was reduced over time following induced internalization of integrin α5. Cell area reduction was <5% in control cells that were also treated with rapamycin, indicating that the reduction is a result of integrin α5 removal specifically, rather than an off-target effect of rapamycin treatment (Figure 2 A,B). Quantification of this change in cell area revealed that after 5 min, there was a decrease of 8% in HeLa, compared with over 20% decrease for SKOV3 cells (Figure 2C, 2D, Supplementary movie 1 and 2). This suggests that SKOV3 cells are more sensitive than HeLa to removal of α5β1 integrin heterodimers from the cell surface.

### Induced internalization of integrin α5 causes fibronectin uptake

Integrin α5β1 binds fibronectin, which raises the question of whether hot-wired endocytosis of this integrin heterodimer can impact the ECM potentially by remodeling fibronectin. To address this question, we introduced pre-labeled fibronectin to the culture medium and induced the internalization of integrin α5 using our system. Confocal imaging revealed the colocalization of labeled fibronectin with α5-mCherry-FRB in cells where internalization of α5 was induced but not in controls where no internalization occurred (Figure 3A). This suggested that extracellular fibronectin could be co-internalized with integrin a5b1 heterodimers upon induction by our system.

**Figure 3.**
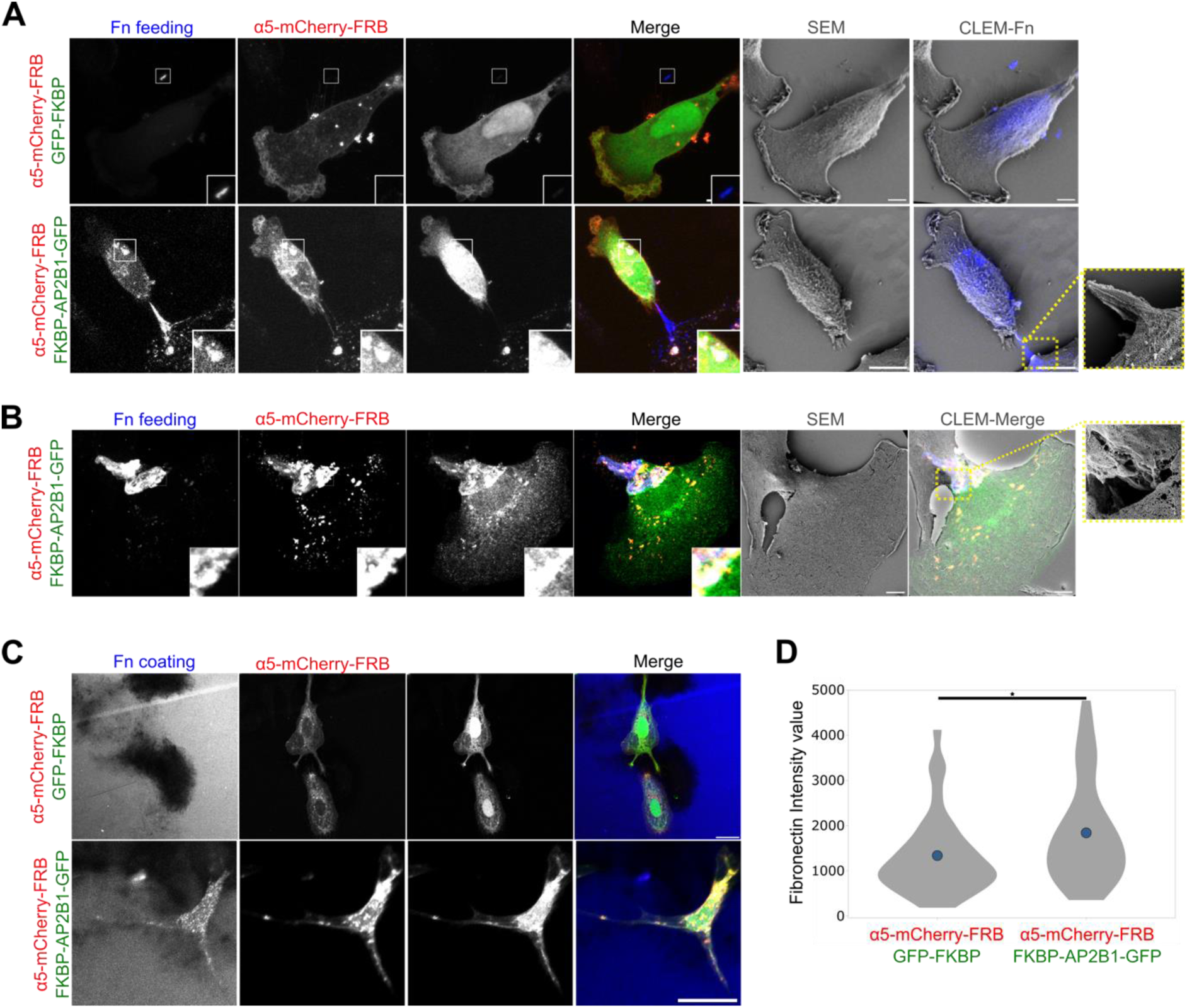
Induced internalization of integrin α5 causes fibronectin uptake. (A) Representative light and electron microscopy images of SKOV3 cells expressing α5-mCherry-FRB (red) and either GFP-FKBP (green, control) or FKBP-AP2B1-GFP (green), treated with rapamycin (200 nM) in the presence of labeled fibronectin (Fn, blue - 20µg/mL). (B) A further example, prepared as in A, showing fibronectin fibers being pulled inside the cells coincident with α5-mCherry-FRB and FKBP-AP2B1-GFP signals. (C) Representative micrographs of cells on labeled fibronectin-coated surfaces (20µg/mL Fn). Increased uptake of fibronectin (blue) is apparent in the cells which have been induced to internalize α5-mCherry-FRB. (D) Violin plots showing the extent of fibronectin uptake per cell. N = 35, p-value<0.05 (GFP-FKBP), 45 (FKBP-AP2B1-GFP), from 3 independent experiments. Scale bars, 10 µm (light and electron microscopy).

To understand the apparent uptake of labeled fibronectin by hot-wiring endocytosis of integrin α5, we applied correlative light and electron microscopy (CLEM). Correlated SEM imaging revealed aggregates of fibronectin atop the cells where α5 internalization had been induced. These aggregates were significantly larger than typically observed, suggesting that the induced internalization of α5 had directly influenced fibronectin clustering, which resulted in pronounced surface aggregation within 30 min (Figure 3A). Using confocal imaging and subsequent 3D projection revealed that the fibronectin was in some cases internal but also that some fibronectin fibers remained external to the cell (Figure 3B, Supplementary S6, supplementary movie 3). Finally, rather than feeding labeled fibronectin to cells we tested if uptake could occur from a supported surface. Cells were cultured on a fibronectin-coated glass surface and the uptake of labelled fibronectin in cells with induced α5 internalization investigated (Figure 3C). Quantification of the total amount of fibronectin per cell revealed ∼30% increase 30 min after induced internalization of integrin α5 compared to control (Figure 3D). Together these results indicate that hot-wired endocytosis of integrin α5 is sufficient to cause the internalization of fibronectin that is presumably associated with the α5β1 integrin being endocytosed.

### Effect of induced α5 internalization on spheroid formation

Does the removal of surface integrin α5β1 heterodimers using hot-wired endocytosis affect cell-cell interactions? To address this question, we studied the effect of induced internalization of α5-mCherry-FRB on the process of spheroid formation. We used honeycomb-well supports (400 µm width, 200 µm height) that were agarose-coated to make them non-adherent. These supports favor the growth of cancer spheroids with reproducibility and size control^24^. SKOV3 cells co-expressing α5-mCherry-FRB and either FKBP-AP2B1-GFP or GFP-FKBP (control) were seeded into honeycomb-wells and rapamycin was added to the wells. Using light microscopy, cells were monitored over 2 days to capture the different stages of spheroid formation. Compared to controls, cells with induced internalization of integrin α5 integrin modulation showed disrupted spheroid formation at 46 h (Figure 4A). At 0 h, the cell coverage of the well is total and upon spheroids formation the fraction of the well covered with cells decreases as the spheroid compacts. We measured the well coverage to be greater and less compacted in cells with induced internalization compared to controls (Figure 4B). These experiments confirm that integrin α5β1 is important for spheroid formation and modulating the surface population of α5 via hot-wired endocytosis is a way to change cell-cell interactions.

**Figure 4.**
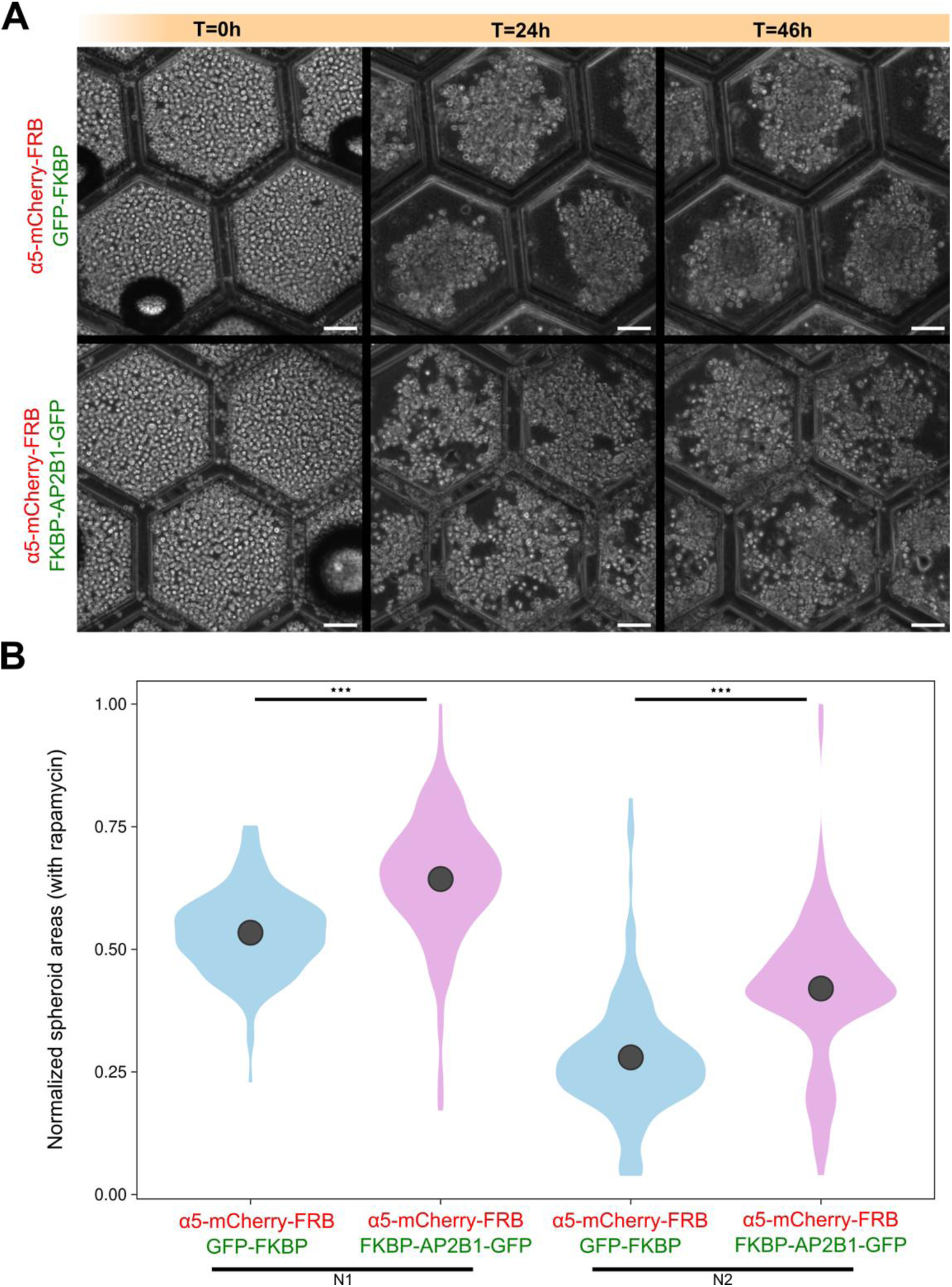
Effect of induced internalization of integrin α5 on spheroid formation. (A) Representative light micrographs of hexagonal wells with SKOV3 cells expressing the indicated constructs and treated with rapamycin. Three time points are shown which capture the compaction of SKOV3 cells into spheroids. Scale bar, 100 µm. (B) Measurement of the total cell area normalized to the well area at 46 h. Over 100 spheroids per condition were analyzed, N1 and N2 are independent replicates, *** p-value<0.001.

## Discussion

In this study, we developed a method to rapidly downregulate the surface population of α5β1 integrins on-demand. As summarized in Figure 5, we showed that targeting a single cell surface receptor in this way leads to rapid cellular responses: altering cell morphology, adhesion dynamics, extracellular matrix remodeling and cell-cell interactions in cancer cells, which are all critical functional factors for cancer progression and metastasis^14,19^.

**Figure 5.**
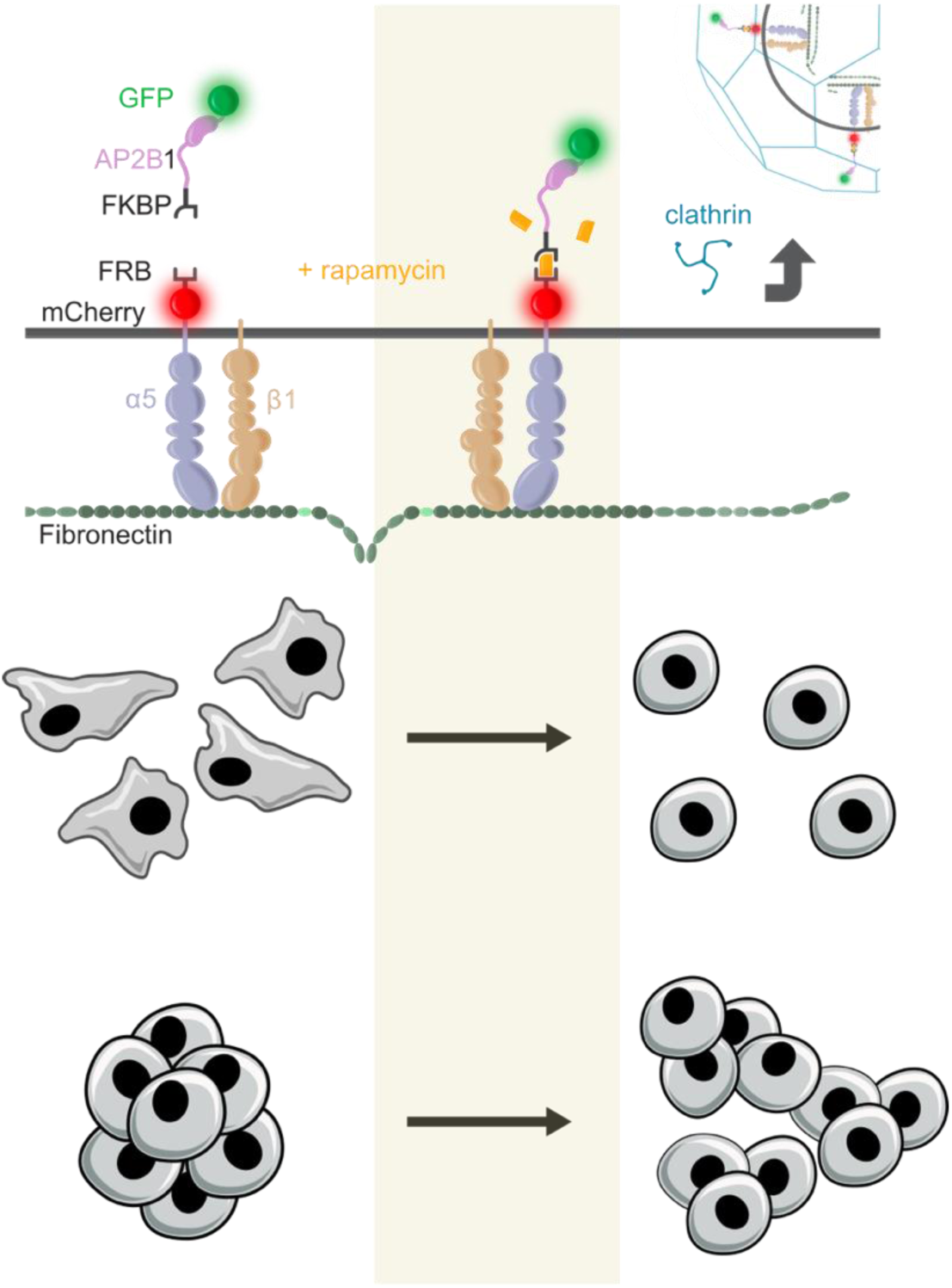
Summary model of rapid tuning of multiscale cell interactions

Rapid modulation of cell surface receptor number is a powerful method to understand their influence on cell biology. Unlike knockdown, knockout or other technologies that affect protein abundance, hot-wired endocytosis allows the down regulation of receptors acutely, on a timescale of minutes. As well as more acute inhibition, it is possible to observe behavior in a cell prior to and following downregulation. For example, we showed that induced α5β1 internalization resulted in a decrease in cell area. Interestingly, a study by Lerche and coworkers, which rapidly increased the surface population of α5β1 using the RUSH system saw the corresponding opposite effect. To do this, they synchronized anterograde transport of integrins from the endoplasmic reticulum, allowing the delivery of new integrins to the plasma membrane, and they reported an increase in cell area, corroborating our findings^25^. More broadly, these results confirm at the single cell level, previous studies suggesting that integrins are not simply adhesion receptors but are pivotal in signaling pathways that influence cell morphology, motility, and survival, as well as cancer processes^1,4,11,12,23,26–30^ .

In addition to influencing cell behavior by downregulating α5β1 at the cell surface, we also found that triggering the removal of α5β1 was sufficient to cause uptake of extracellular fibronectin. The amount of uptake was surprisingly large and have not been previously reported in ovarian cancer cells. Therefore, this method has the potential application for ECM remodeling (Bascetin et al., 2018)^31^ or, when alternative receptors are targeted, for inducible uptake of other large molecules.

Integrin α5β1 heterodimers have long been considered as a key molecular partner for spheroid formation^14^. Induced internalization of α5β1 not only altered cell-ECM interactions but also cell-cell interactions as evidenced by disrupting spheroid formation with SKOV3 cells. This disruption is in agreement with previous work, but challenges the conventional understanding of spheroid stability and highlights the potential of targeting integrins in disrupting tumor cell aggregates.

The rapid response/readout of our system enables the exploration of the immediate cellular responses to changes in integrin availability, which is crucial for understanding the integrin signaling dynamics that underly rapid cellular adaptations to changing environments.

In conclusion, the development and application of the FKBP-rapamycin-FRB system to study integrin dynamics represent a powerful approach for elucidating the complex roles of integrins in cellular and pathological processes. This study not only advances our understanding of integrin function but promises to enhance our comprehension of integrin-mediated cellular processes in health and disease, traversing a wide array of biological complexities and offering significant implications for the development of targeted cancer therapies.

## Materials and Methods

### Cell Culture

The Human ovarian adenocarcinoma cell line, SKOV3 (ATCC HTB-77), was cultured in RPMI-1640 GlutaMAX (Gibco Ref-72400021) supplemented with 10% fetal bovine serum (FBS) (Biowest S181G-500), 7.5% sodium bicarbonate, penicillin (200 IU/mL), and streptomycin (200 µg/mL). Human ovarian adenocarcinoma cell line, NIH:OVCAR-3 (ATCC HTB-161), was also cultured under similar conditions using RPMI-1640 GlutaMAX supplemented with 10% fetal bovine serum (FBS), 7.5% sodium bicarbonate, penicillin (200 IU/mL), and streptomycin (200 µg/mL). HeLa cells (Health Protection Agency/European Collection of Authenticated Cell Cultures 93021013; Research Resource Identifier [RRID]: CVCL_0030) were cultured in DMEM (Dulbecco’s Modified Eagle Medium) supplemented with 10% fetal bovine serum (FBS), penicillin (200 IU/mL), and streptomycin (200 µg/mL).

All cell lines were maintained in a humidified incubator at 37°C with 5% CO₂ and confirmed to be mycoplasma-free by routine PCR-based testing.

For transfection, cells were seeded 24 hours prior to reach 70-80% confluency. SKOV3 cells were transfected using X-tremeGENE HP transfection reagent (Roche, XTGHP-RO) according to the manufacturer’s instructions. Cells were typically analyzed 48 hours post-transfection.

### Molecular biology

To generate α5-mCherry-FRB, mCherry-FRB was amplified from CD8-mCherry-FRB (Addgene #100738) and inserted into Alpha5 integrin-GFP (Addgene #15238) using SacII and MfeI sites, replacing EGFP. PCR was with the following primers: - Forward: ATTAATCCGCGGTGGTCGCCACCATGGTGAG; Reverse: CCGCGTTAACAACAACAATTGCATTC. Plasmids to express GFP-FKBP or FKBP-AP2B1-GFP (Addgene #100726) were available from previous work (Wood et al., 2017)^20^.

### Immunofluorescence

Cells were fixed with 4% paraformaldehyde (PFA) for 15-20 min at room temperature (RT). After fixation, cells were washed three times in phosphate-buffered saline (PBS) and incubated with 0.5% bovine serum albumin (BSA) in PBS for 30 minutes at RT to block non-specific binding. Cells were then permeabilized with 0.1% Triton-X in PBS for 5 min at 4°C.

Primary antibody solutions were prepared in PBS with 0.5% BSA, and cells were incubated with the primary antibody solutions for 2 h at RT. The primary antibodies used included mouse anti-Integrin alpha 5/CD49e (Snaka51, Catalog Number: MABT201, Sigma-Aldrich, Dilution: 1:500), mouse anti-Integrin alpha V (L230, Catalog Number: L230, Enzo Life Sciences, Dilution: 1:100), mouse anti-EEA1 (Catalog Number: 610457, BD Biosciences, Dilution: 1:200), and rabbit anti-Fibronectin (Catalog Number: F3648, Sigma-Aldrich, Dilution: 1:50). After primary antibody incubation, coverslips were washed five times in PBS with 0.5% BSA at RT.

Secondary antibodies solutions were prepared in PBS, and cells were incubated with the secondary antibodies solutions for 1 hour at RT. The secondary antibodies used included anti-rabbit Alexa Fluor 488 (Catalog Number: A-11008, Invitrogen, Dilution: 1:200), anti-rabbit Alexa Fluor 568 (Invitrogen, Dilution: 1:200), anti-mouse Alexa Fluor 488 (Catalog Number: A-11029, Invitrogen, Dilution: 1:200), and anti-rabbit CF640R (Sigma-Aldrich, Dilution: 1:100). Following secondary antibody incubation, cells were washed with PBS and before mounting on glass slides using Prolong Diamond or Prolong Gold mounting media and sealed with dental silicone (Picodent).

### Labeling of Fibronectin

Fibronectin was isolated from human blood plasma using an established protocol^32^. Briefly, 1.5-2 mL of purified fibronectin at a concentration of 2 mg/mL was dialyzed twice in 0.1 M carbonate/bicarbonate buffer at pH 9.2. Subsequently, a mixture comprising 5 µL of Alexa fluorophore 405 at a concentration of 10 mg/mL in DMSO was homogenized with 1 mL of fibronectin solution and incubated for 1 h at RT in the dark. To separate the dye-bound fibronectin from free dyes, gel filtration chromatography was used (Sephadex G25 SIGMA G251250). Fractions were collected and analyzed by spectrophotometry using wavelengths of 280 and 405 nm. Corrected absorbance values of fibronectin-Alexa Fluor were computed utilizing the formula A_prot_ = A_280_ – A_405/Alexa_ x CF, where CF represents a correction factor. Prior to use in cell culture experiments, all samples were filtered to ensure sterility.

### Microscopy and analysis

The microscopy data were acquired using two systems: Confocal laser scanning microscope (Zeiss LSM 900) equipped with a 405 nm diode laser, 458, 488, 514 nm argon lasers, and 561, 633 nm HeNe lasers. The objectives in the microscope include 63× (1.4 NA, oil). All the imaging parameters were set using the Zen software according to Nyquist criterion (raw data shared). Nikon SoRA spinning disk microscope. The 100× (oil) objective was used for all acquisitions and time-lapse observations. NiS-Elements software was used to set the parameters and image acquisition (see raw data, metadata).

Analysis of data was done using ImageJ/Fiji and analysis pipelines were developed in Python (Boché et al., 2024)^33^.

### CLEM

Cells were cultured on gridded glass bottom petri dishes (Mattek, USA) and fixed with 4% PFA in PBS for 5 min. After washing, cells were left in PBS and imaged by confocal microscopy. Coverslips were then fixed with a mixture of PFA 2%, glutaraldehyde 2.5% in 0.2 M sodium cacodylate, pH 7.5, washed with cacodylate buffer, and dehydrated through graded ethanol series from 30% to 100%. Then, cells were critical point dried (CPD300, Leica) and finally coverslips were mounted on SEM stubs with carbon tape and metallized with 4 nm of platinum (Leica ACE600). Electron microscopy was carried out using a field emission gun scanning electron microscope (SEM, GeminiSEM300, Carl Zeiss) with an acceleration voltage of 2 keV under high vacuum. Secondary electrons were collected. Scan speed and line averaging were adjusted during observation. Images were correlated and were processed with Imagej/Fiji software and aligned with ICY software using the ecCLEM plugin^34^.

### Spheroid culture

Microfabricated patches with a honeycomb pattern were used for cell culture, facilitating mechanical and biological exchanges crucial for cell maintenance. These patches were fabricated using photolithography and soft-lithography techniques, followed by gold-coating and layering with electrospun gelatin and agarose to promote non-adherent conditions, as described by Changchong et al. (2022)^24^. SKOV-3 cells were seeded on these patches, allowing for controlled spheroid size and homogeneity through the adjustment of microframe dimensions and cell seeding density, ensuring uniform and isolated spheroids across all compartments. Patches with wells measuring 400 x 200 µm (w x h) were used to seed SKOV-3 cells in normal culture media. Polydimethylsiloxane (PDMS) molds were used to secure the patches within a well of a 12-well plate. Spheroid growth was monitored via time-lapse recordings using the Leica Thunder Live Cell Imager, under controlled conditions of a 37°C humidified atmosphere containing 5% CO₂. The area spread of spheroids was measured over time, and normalized spheroid areas were plotted to assess growth patterns.

## ACKNOWLEDGMENTS

The authors acknowledge the imaging facility “Microscopies&Analyses” of CY Cergy Paris Université where confocal and electron microscopy were carried out. We would like to thank the Conseil regional Île de France for funding the COMICER project ( SESAME grant: 19200103) that has allowed the acquisition of a ZEISS LSM710, the CERASEM project (SESAME grant:15013107) for the acquisition of a ZEISS Gemini SEM 300 and the COMCOI@CY project (SESAME grant: 00082089) for the Zeiss LSM900. SK was supported by a EUTOPIA Co-Tutelle PhD Studentship. The work was supported by an HFSP Research Grant to SJR (RGP0025/2022) and Agence Nationale de la recherche, ANR JCJC Modulo-EMT (Projet-ANR-21-CE19-0006). We thank Gabrielle Larocque and Adrien Assié for her assistance with schematic diagrams, and also Maelle Lorvellec and Laura Cooper from Warwick Computing and Advanced Microscopy Unit (CAMDU) for their help and support.

## Supplementary Information

**Figure S1:**
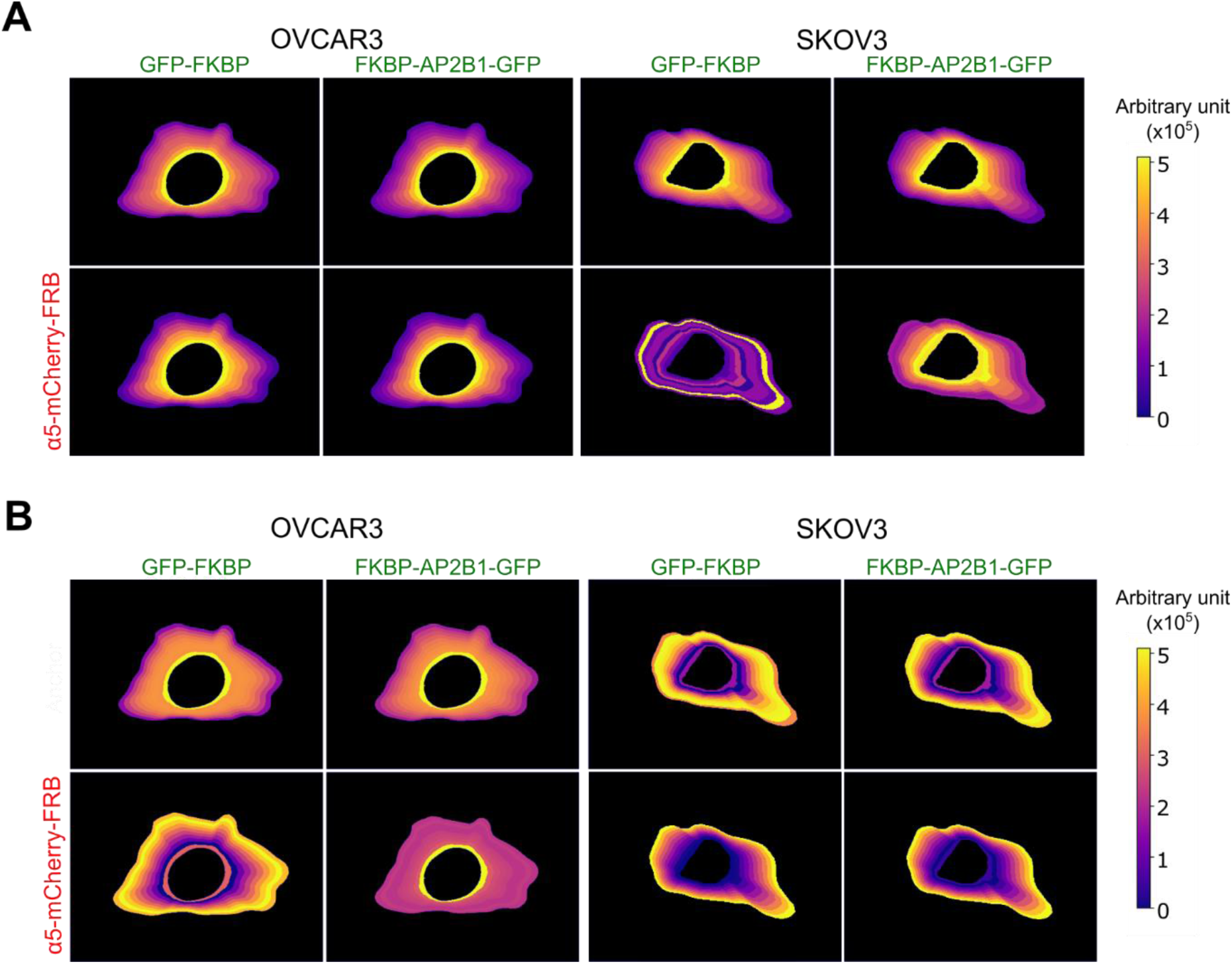
Distribution of α5-mCherry-FRB and GFP-FKBP or FKBP-AP2B1-GFP as a function of distance from nucleus to the membrane after cell segmentation. (A) Analysis of max intensity fluorescence per section for SKOV3 and OVCAR3. (B) Analysis of fluorescence raw integrated density per section for SKOV3 and OVCAR3 for >150 cells per condition in 3 independent experiments.

**Figure S2:**
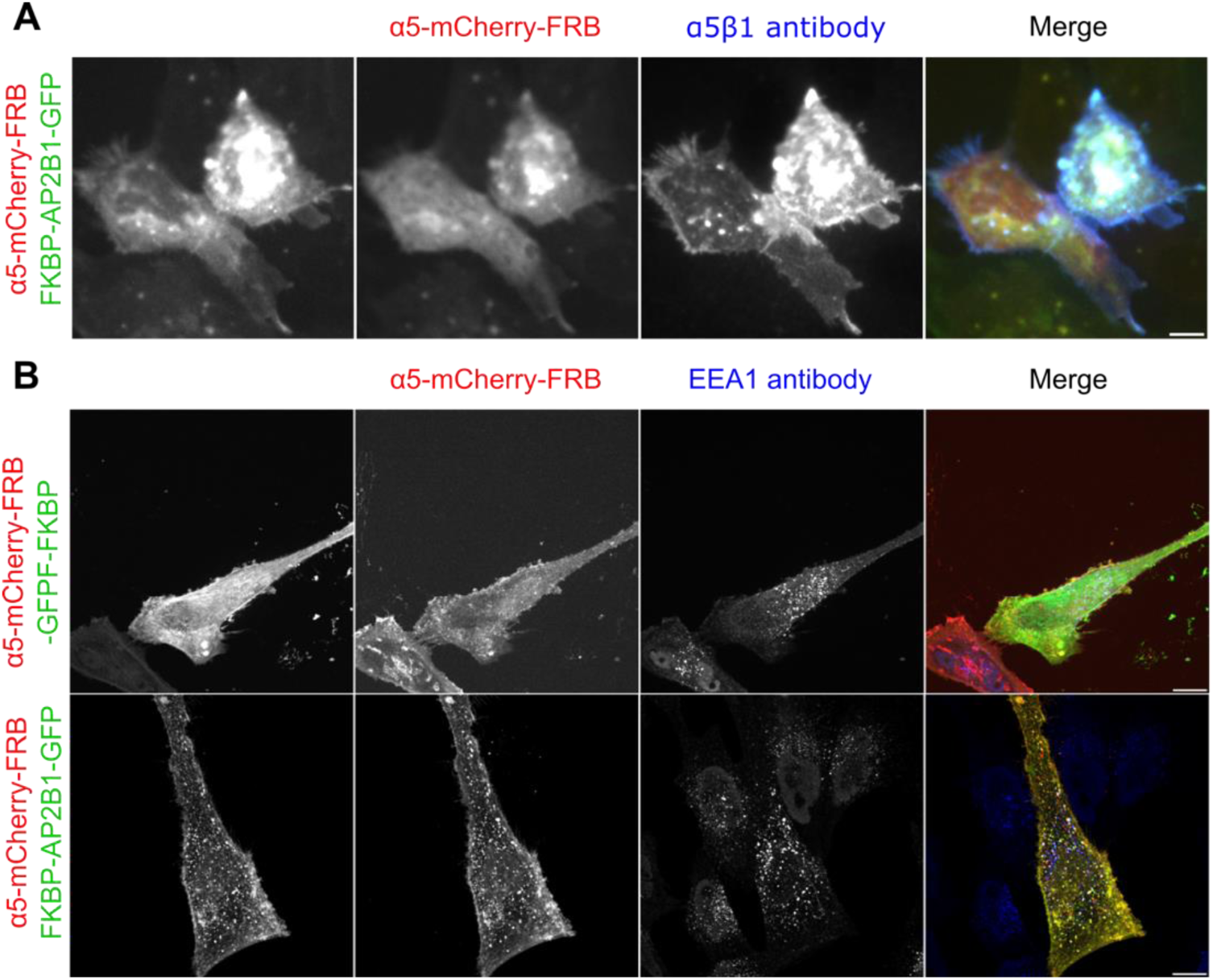
Induced internalization of α5 integrins. A) HeLa cells observed with anti-α5β1 antibody feeding over 15 minutes followed by washes, The internalized α5 vesicles colocalize with anti-α5β1 antibody B) HeLa cells observed with EEA1 antibody staining. The internalized α5 vesicles colocalize with Early endosomal marker EEA1, showcasing the induced endocytosis α5 vesicles follow normal endocytic pathway.

**Figure S3:**
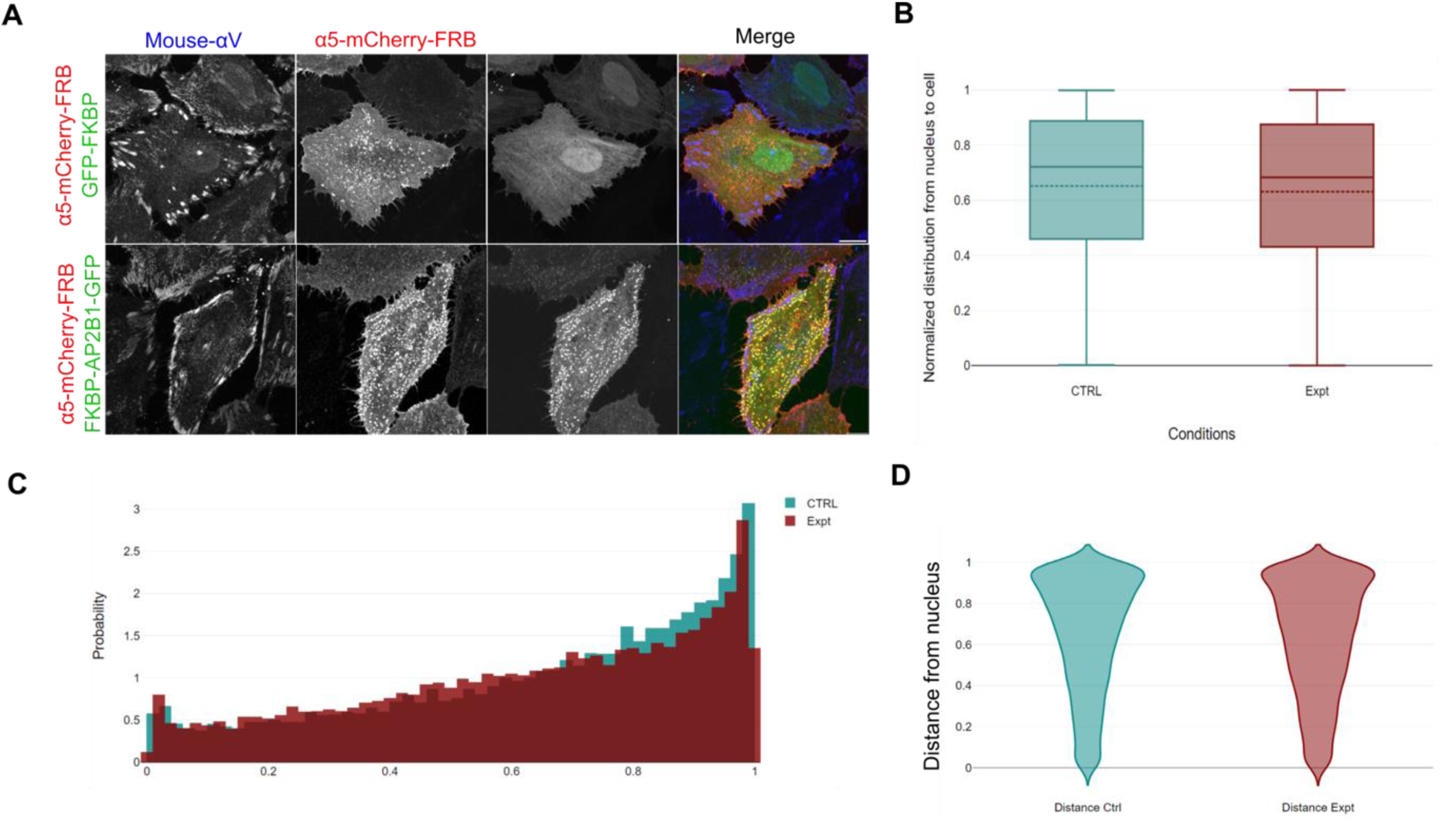
Induced internalization of α5 integrins has no effect on αV localization. (A) Representative micrographs of HeLa cells expressing the indicated constructs, treated with rapamycin and detected with anti-αV (blue). Scale bar, 10 µm. (B-D) Quantification of αV signal in control cells and those with induced internalization of α5 in HeLa cells.

**Supplement S4.**
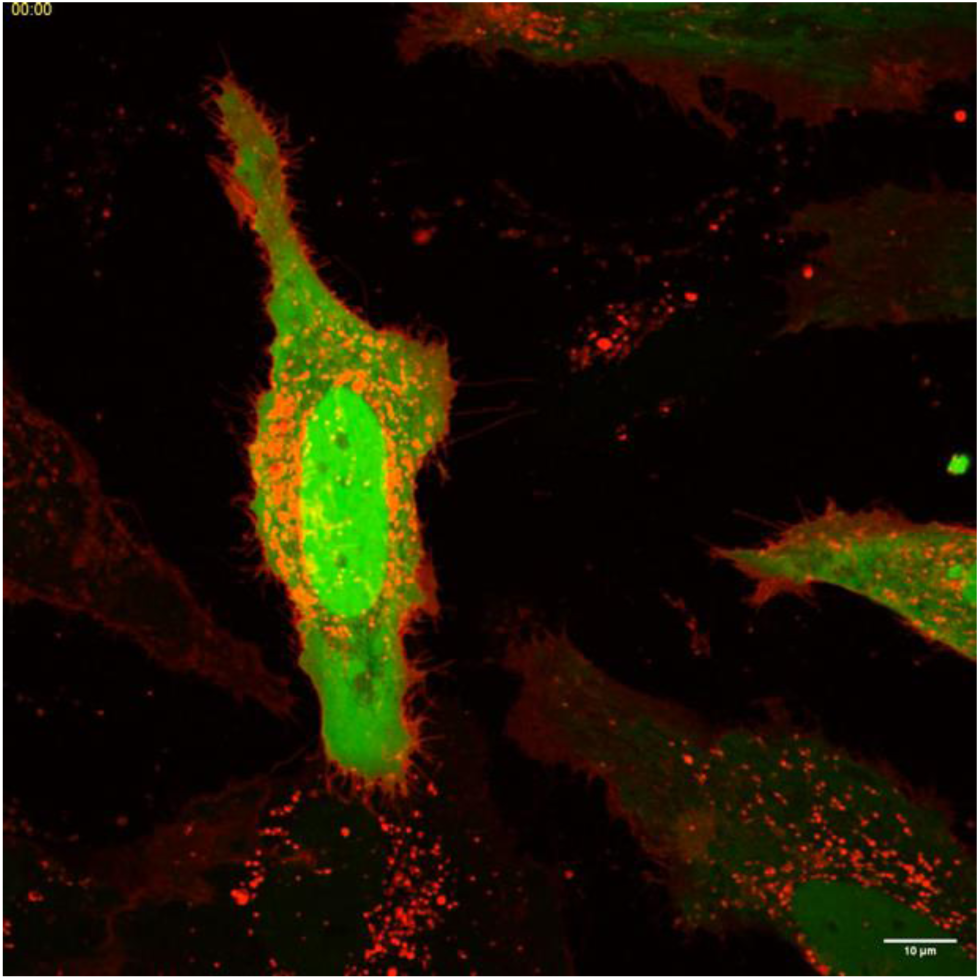
Live cell imaging of control HeLa cells expressing α5-mCherry-FRB (red) and GFP-FKBP (green), treated with rapamycin (200 nM). Movie, 1 minute per frame for 30 mins.

**Supplement S5.**
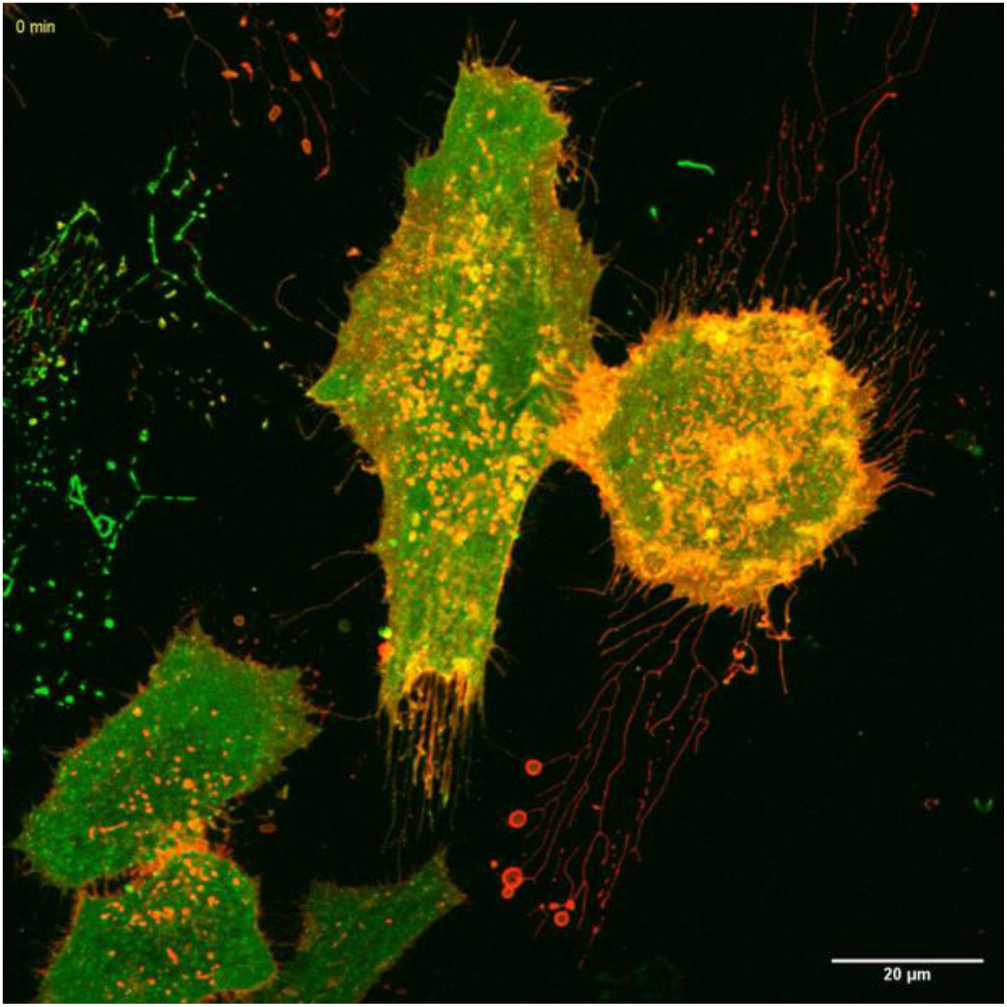
Live cell imaging of HeLa cells expressing α5-mCherry-FRB (red) and FKBP-AP2B1-GFP (green), treated with rapamycin (200 nM). Movie, 1 minute per frame for 30 mins. Note the formation of puncta and the cell area decrease.

**Supplement S6:**
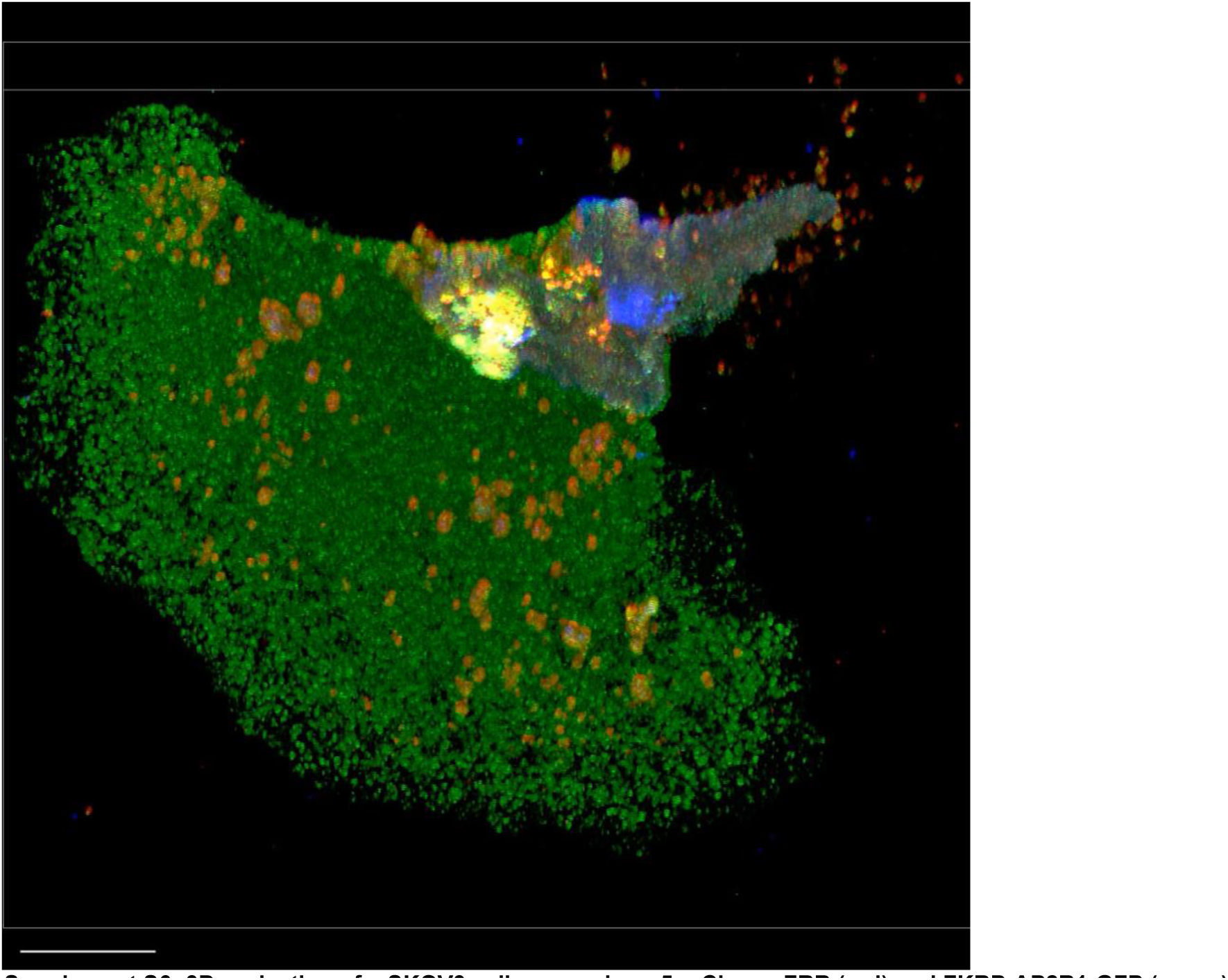
3D projection of a SKOV3 cell expressing α5-mCherry-FRB (red) and FKBP-AP2B1-GFP (green) showing uptake of fibronectin (blue). Cells were fed labeled fibronectin along with rapamycin to induce internalization of α5-mCherry-FRB when the internalization was triggered. Scale bar, 10 µm.

